# Inflammatory responses following CRISPR modification of the nuclear localisation sequence in endogenous interleukin-1α

**DOI:** 10.1101/2025.09.30.679460

**Authors:** Christopher Hoyle, Rodrigo Díaz Pino, Si Min Lai, Jack P Green, Antony Adamson, Graham Coutts, Catherine B Lawrence, Mark A Travis, David Brough, Gloria Lopez-Castejon

## Abstract

Interleukin (IL)-1α is a pro-inflammatory member of the IL-1 cytokine superfamily and is important for inflammatory responses to infection and injury. Unlike pro-IL-1β, pro-IL-1α is mainly localised to the nucleus upon expression. This is mediated by a nuclear localisation sequence (NLS) responsible for its importin-dependent transport into the nucleus. This nuclear localisation and the presence of histone acetyl transferase (HAT)-binding domains within the pro-domain suggest a role of this cytokine in gene transcription regulation. In addition, nuclear trafficking of pro-IL-1α is proposed to regulate its secretion. To-date, studies on the nuclear role of pro-IL-1α have used overexpression systems. Here, we generated a mouse where the endogenous *Il1a* gene was edited with CRISPR to disrupt the NLS (mNLS). Using an *in vitro* approach with murine macrophages we found that this NLS mutation did not affect pro-IL-1α RNA expression levels in response to LPS but increased its protein expression levels. Moreover, we found that the transcriptional signature induced by LPS was not altered between WT and mNLS macrophages. Release of IL-1α in response to different stimuli such as ionomycin was not negatively impacted by disrupted nuclear localisation, although higher levels of IL-1α release were detected, potentially due to increased levels of pro-IL-1α. Inflammatory responses in an *in vivo* model of peritonitis and an influenza infection model were comparable between WT and mNLS mice. Thus, we have established a mouse model in which pro-IL-1α nuclear localisation is disrupted, although future research is required to reveal the importance of this nuclear localisation for IL-1α function.

## Introduction

Interleukin-1α (IL-1α) is a pro-inflammatory member of the IL-1 cytokine superfamily and is important for inflammatory responses to infection and injury (1). IL-1α is produced as a 31kDa precursor called pro-IL-1α, proteolytic cleavage of which produces a 17kDa mature protein of increased biological activity that is released and signals via the type 1 IL-1 receptor (IL-1R1) on neighbouring cells (1). IL-1α is closely related to IL-1β, another proinflammatory member of the IL-1 family, and from which it arose as a gene duplication in mammals (2). Whilst both IL-1α and IL-1β share a receptor through which they can initiate inflammatory signalling (IL-1R1), some additional functionality is attributed to pro-IL-1α due to the presence of a nuclear localisation sequence (NLS) and histone acetyl transferase (HAT)-binding domains within the pro-domain (2-4). In general, the pro-domain of pro-IL-1α and its NLS is well conserved amongst most mammals although is lost in a number of species (2, 5). When intact, the NLS on pro-IL-1α is responsible for its importin-dependent transport into the nucleus (6, 7).

In addition to nuclear actions, we have also hypothesised previously that nuclear trafficking of pro-IL-1α could act to regulate its secretion. Unlike pro-IL-1β, pro-IL-1α is not directly activated by caspase-1 following inflammasome activation. Cleavage depends upon calcium-activated calpain proteases (8) that become active in response to stimuli that increase intracellular calcium concentration including inflammasome activators like extracellular ATP although other proteases have also been described (9, 10). Once processed, mature IL-1α is secreted via unconventional protein secretion pathways (8) or is released by cell death (11). We have previously reported that in COS7 cells transfected to express either pro-IL-1α-GFP or GFP alone, the release of IL-1α was limited by its retention in the nucleus (11). Furthermore, in HeLa cells transiently transfected to express wild type (WT) pro-IL-1α, or pro-IL-1α in which the NLS was mutated to lose function and hence mainly present in the cytosol, more IL-1α was secreted from cells expressing the mutant in response to IL-1α secretion stimuli such as the calcium ionophore ionomycin (5).

IL-1α is important for an effective anti-viral host-response, particularly where the virus has host evasion mechanisms. HSV-1 infection inhibits the release of IL-1β via an immune evasion mechanism and IL-1α is essential for a protective immune response (12). Orzalli et al. established that in response to viruses that can evade PRR-dependent anti-viral gene expression, the IL-1 cytokines IL-1α and IL-1β provide a back-up anti-viral system where these cytokines induce expression of anti-viral genes and limit viral replication (13). In a follow-on study, Orzalli et al. suggest that in human keratinocytes, viral disruption of host cell protein synthesis activates guard proteins that leads to a gasdermin E-dependent pyroptosis and IL-1α release (14). Thus, losing the NLS could benefit an organism fighting viral infection by increasing levels of released IL-1α. This was our rationale for the following study, where we aimed to establish the importance of nuclear trafficking during inflammation and its potential importance for effective antiviral inflammatory responses.

## Results

### Mutation of pro-IL-1α NLS leads to loss of nuclear localisation

Our previous studies on mutating the NLS to investigate its effect on secretion of IL-1α relied upon ectopic expression systems (5, 11). Here we wanted to study the effect of nuclear trafficking on endogenous IL-1α and to do this we employed CRISPR to mutate the NLS. In our first attempt we generated mice with mutations in the NLS using a CRISPR strategy that resulted in seven base substitutions (15). These substitutions were combinations of the desired mutations and some synonymous, silent mutations intended to prevent re-editing of the allele and make detectable changes for follow on colony genotyping. However, analysis of these mice showed a loss of *Il1a* gene expression. Further analysis of the gene locus indicated that the combination of substitutions likely impacted the sequence and structure of a lncRNA gene found antisense to *Il1a* (now named AS-IL1α), which has cis-regulatory functions for *Il1a* itself (16). To prevent this collateral gene perturbation, we designed a second strategy to minimise the number of base substitutions generated, creating just two base changes in the NLS encoding residues. We used RNAsnp to predict the impact of these changes on AS-IL1α structure, which indicated no significant structural change. The two base substitutions mutated a critical motif within the NLS of the endogenous pro-IL-1α (^85^KKRR^88^) to the non-functional NLS sequence shared by almost all toothed whales (^85^KNRW^88^) that we had previously shown to prevent IL-1α nuclear localisation in an ectopic expression system (5) (Figure 1A). These mutant mice were henceforth called mutant NLS, or mNLS.

**Figure 1.**
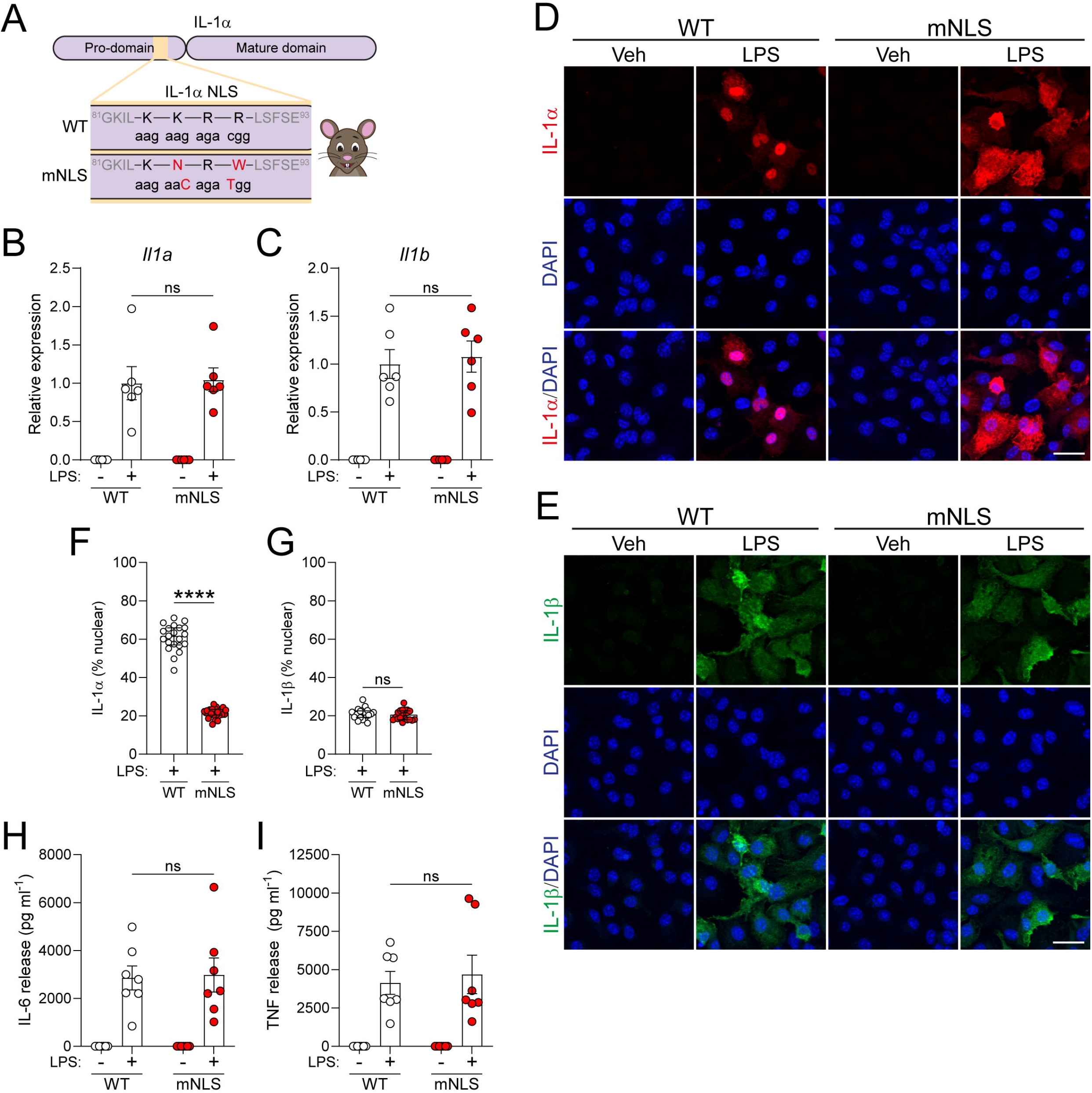
Pro-IL-1α NLS mutation does not affect expression but reduces nuclear localisation. (**A**) Schematic of pro-IL-1α NLS mutation in the mNLS mouse. (**B-I**) WT or mNLS BMDMs were primed with vehicle (PBS) or LPS (1 µg/ml, 4 h). qPCR analysis of (B) *Il1a* and (C) *Il1b* expression (n=6). Immunofluorescence labelling of (D) pro-IL-1α and (E) pro-IL-1β, and quantification of nuclear localisation of (F) pro-IL-1α and (G) pro-IL-1β (n=7). Scale bar is 20 µm. Release of (H) IL-6 and (I) TNF into the supernatant (n=7). Data are presented as mean ± SEM (B,C,H,I), or median ± IQR (F,G). Data were analysed using unpaired t-test (C,F,G,H) or Mann-Whitney test (B,I). ****P<0.0001; ns, not significant.

Bone marrow-derived macrophages (BMDMs) were isolated and cultured from WT and mNLS mice, treated with or without LPS and subsequently analysed by qPCR. We confirmed that both IL-1α and IL-1β were expressed, and that expression levels were similar between WT and mNLS BMDMs (Figure 1B, C). A similar experiment was conducted but the cells were fixed and immunocytochemistry was used to analyse pro-IL-1α and pro-IL-1β expression and cellular localisation. In WT mice, pro-IL-1α was largely located in the nucleus, as expected, while pro-IL-1β was diffusely present throughout the cell (Figure 1D-G). Strikingly, in mNLS mice the pro-IL-1α was no longer concentrated in the nucleus but was diffusely distributed throughout the cell, similar to pro-IL-1β (Figure 1D-G). LPS-induced IL-6 and TNF release was similar between the WT and mNLS BMDMs, indicating that TLR4 signalling was unaffected (Figure 1H, I). We also examined pro-IL-1α subcellular localisation in peritoneal macrophages isolated from these mice and subsequently primed with LPS, and detected nuclear pro-IL-1α in WT mice and cytosolic pro-IL-1α in mNLS mice (Supplementary Fig 1).These data show that our mNLS mice express normal levels of pro-IL-1α, and that the mutations of the NLS had disrupted nuclear localisation of pro-IL-1α.

As nuclear IL-1α is proposed to influence transcriptional regulation (17), we next performed bulk RNA-Sequencing in BMDMs treated with vehicle or LPS. We observed that LPS induced a large transcriptional response in the WT and mNLS BMDMs (Fig 2A-C), however there were no differences in this response between the WT and mNLS cells (Figure 2D-F).

**Figure 2.**
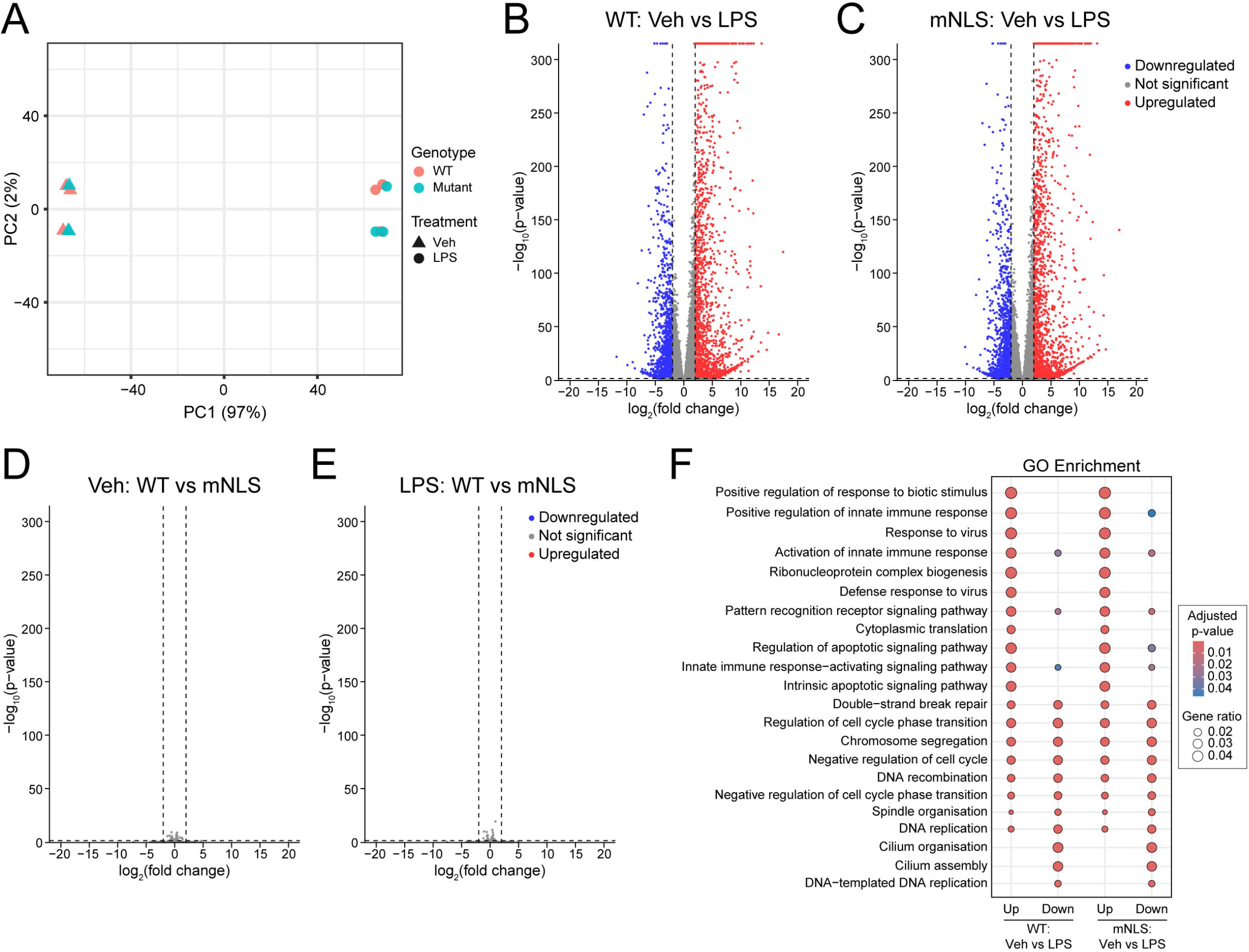
mNLS IL-1α does not affect LPS-induced gene expression in BMDMs. WT or mNLS BMDMs were primed with vehicle (PBS) or LPS (1 µg/ml, 4 h) (n = 3), followed by bulk RNA-Seq analysis. **(A)** Principal component analysis (PCA) of RNA-Seq data showing clustering by genotype and treatment. **(B)** Volcano plot of differentially expressed genes in WT BMDMs (Veh versus LPS). **(C)** Volcano plot of differentially expressed genes in mNLS BMDMs (Veh versus LPS). **(D)** Volcano plot of differentially expressed genes between WT and mNLS BMDMs after vehicle treatment. **(E)** Volcano plot of differentially expressed genes between WT and mNLS BMDMs after LPS treatment. **(F)** Gene ontology (GO) enrichment analysis of biological processes for differentially expressed genes in WT (Veh versus LPS, left) and mNLS (Veh versus LPS, right) BMDMs.

### Loss of nuclear localisation does not impair pro-IL-1α processing and release

Having shown that mNLS IL-1α did not affect LPS-induced gene expression, we set out to determine whether disrupted nuclear localisation of pro-IL-1α affected its processing and release. BMDMs were treated with LPS to induce IL-1α expression and then treated with ionomycin to induce calpain-dependent processing and release of IL-1α. Some cells were also pre-treated with MCC950, an NLRP3 inflammasome inhibitor, or calpeptin, a calpain inhibitor that prevents IL-1α cleavage. mNLS BMDMs exhibited robust IL-1α release in response to ionomycin, and this was higher than in WT BMDMs, while levels of released IL-1β and LDH release, an indicator of cell death, were unaffected between WT and mNLS cells (Figure 3A-C). MCC950 did not affect IL-1α cleavage or release, whereas calpeptin strongly reduced IL-1α cleavage, although it did not prevent ionomycin-induced cell death, and thus we still detected passive release of pro-IL-1α into the supernatant which could be detected by ELISA and western blotting (Figure 3A, C, D). The IL-1β released into the supernatant was almost exclusively pro-IL-1β in response to cell death (Figure 3D, Supplementary Figure 2), indicating a lack of NLRP3 inflammasome activation in response to ionomycin, as expected. This further explains why MCC950 did not block IL-1β release in response to ionomycin. Levels of pro-IL-1α protein in the cell lysates were higher in the mNLS BMDMs in response to LPS priming, potentially contributing to the observed enhanced levels of IL-1α cleavage and release (Figure 3D, Supplementary Figure 2).

**Figure 3.**
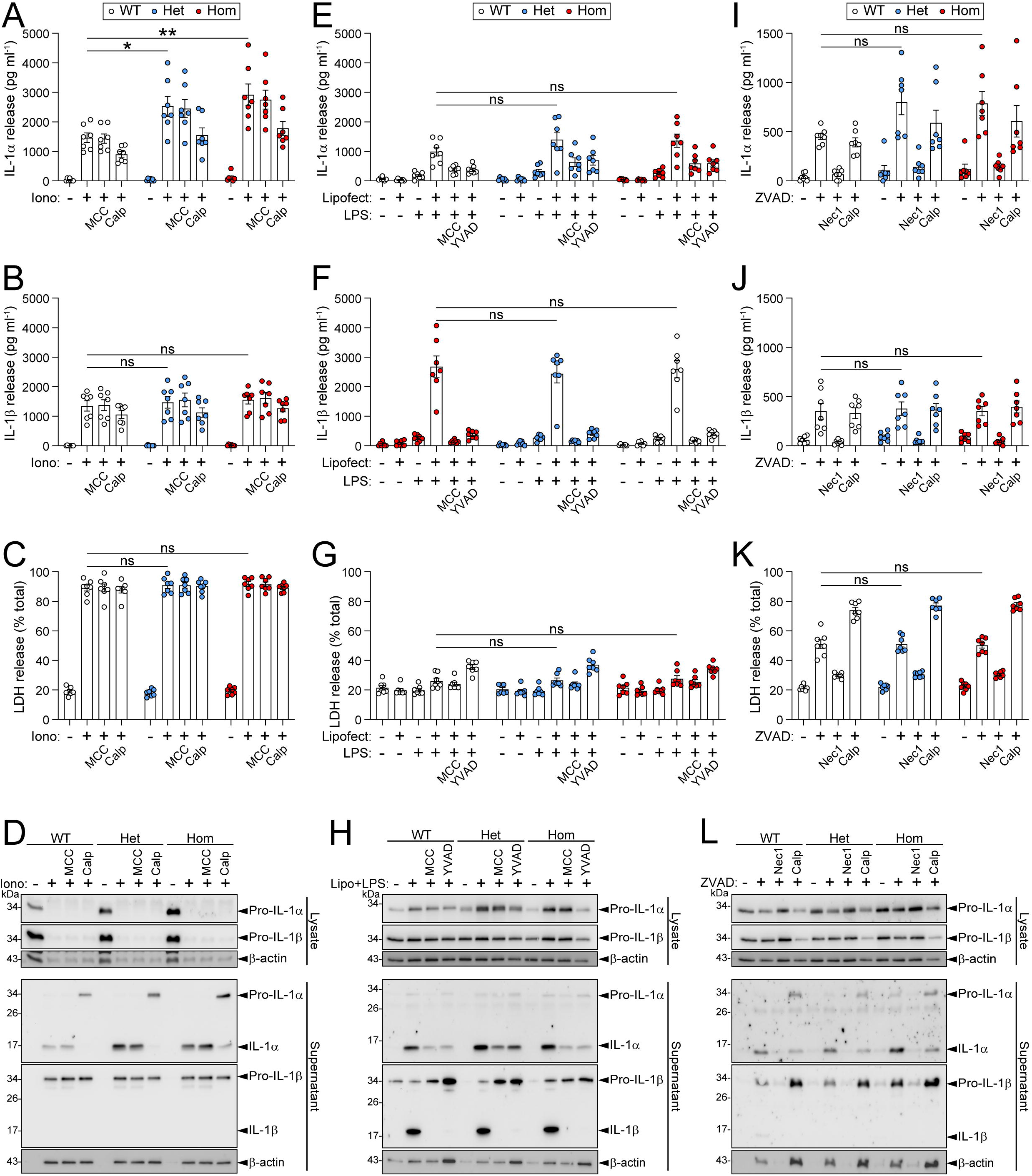
Pro-IL-1α NLS mutation does not negatively affect processing and release of IL-1α. **(A-D)** WT or mNLS BMDMs were primed with LPS (1 µg/ml, 4 h), followed by ionomycin treatment (10 µM, 1 h) in the presence or absence of MCC950 (10 µM; MCC) or calpeptin (40 µM; Calp). Supernatants were assessed for (A) IL-1α release, (B) IL-1β release, and (C) LDH release. (D) Cell lysates and supernatants were assessed for IL-1α and IL-1β expression and processing by western blotting. **(E-H)** WT or mNLS BMDMs were primed with Pam3CSK4 (100 ng/ml, 4 h), before transfection of LPS (2 µg/ml, 18 h) in the presence or absence of MCC950 (10 μM) or YVAD (100 μM). Supernatants were assessed for (E) IL-1α release, (F) IL-1β release, and (G) LDH release. (H) Cell lysates and supernatants were assessed for IL-1α and IL-1β expression and processing by western blotting. **(I-L)** WT or mNLS BMDMs were primed with LPS (1 µg/ml, 4 h), followed by ZVAD treatment (50 µM, 5 h) in the presence or absence of necrostatin-1 (50 μM; Nec1) or calpeptin (40 μM). Supernatants were assessed for (I) IL-1α release, (J) IL-1β release, and (K) LDH release. (L) Cell lysates and supernatants were assessed for IL-1α and IL-1β expression and processing by western blotting. Data are presented as mean ± SEM. Data were analysed using one-way ANOVA followed by Dunnett’s post-hoc test (A,B,C,E,F,I,J,K) or Kruskal-Wallis test followed by Dunn’s post-hoc test (G). *P<0.05; *P<0.01; ns, not significant.

IL-1α is also reported to be released from macrophages in response to canonical NLRP3 inflammasome activation (18), non-canonical inflammasome activation (9), and in response to necroptosis (19). Canonical NLRP3 inflammasome activation triggers Ca^2+^ entry through gasdermin D membrane pores that activates calpain-dependent IL-1α processing and release (20). In response to LPS treatment followed by the canonical NLRP3 activators nigericin and ATP, IL-1α was processed and released from mNLS BMDMs, and this release was modestly increased compared to WT BMDMs although it was not statistically significant (Supplementary Figure 3A, D). The NLRP3 inhibitor MCC950 blocked IL-1α release in response to nigericin, and it also blocked IL-1β release and cell death as expected, suggesting that IL-1α release was occurring downstream of NLRP3 inflammasome activation (Supplementary Figure 3A-C). Calpain inhibition via calpeptin treatment did not prevent IL-1α release in response to nigericin, as cell death was not prevented and thus this was likely to be passive release of pro-IL-1α, similar to ionomycin treatment (Supplementary Figure 3A, C). In response to ATP, MCC950 treatment did not prevent ATP-induced cell death, hence some IL-1α release was observed, whereas calpeptin caused a greater reduction in IL-1α release, and again this was likely to be unprocessed pro-IL-1α (Supplementary Figure 3D). IL-1β release induced by nigericin or ATP was consistent between WT and mNLS BMDMs, and was blocked by NLRP3 inhibition (Supplementary Figure 3B, E). Activation of the non-canonical inflammasome, stimulated by transfection with LPS, caused comparable IL-1α and IL-1β processing and release from WT and mNLS BMDMs, with little cell death detected (Figure 3E-H, Supplementary Figure 4A-F). Both MCC950 and YVAD, a caspase-1 inhibitor, reduced IL-1α cleavage and release, and strongly reduced IL-1β release, indicating that these events required NLRP3 and caspase-1 activity. The induction of necroptosis in response to LPS and the pan-caspase inhibitor ZVAD also induced IL-1α release from mNLS BMDMs that was modestly increased compared to WT, with again, levels of IL-1β release remaining the same between the cells (Figure 3I-L, Supplementary Figure 4G-L). The RIPK1 inhibitor necrostatin-1 blocked IL-1α and IL-1β release, as well as cell death, in response to ZVAD, confirming that these events were occurring due to necroptosis. Calpeptin reduced IL-1α cleavage in response to ZVAD, but did not prevent release (Figure 3I, L). Together, these data suggest that the processing and release of pro-IL-1α does not depend upon its nuclear localisation, as the loss of pro-IL-1α nuclear localisation did not negatively impact its processing and release. It also appears that loss of nuclear localisation may lead to higher pro-IL-1α protein levels, and hence increased IL-1α release, suggesting that nuclear localisation may thus limit IL-1α release. This is consistent with previously published data utilising ectopic over expression systems, and that in species where the NLS is lost, one may predict greater levels of IL-1α release during inflammation.

### mNLS mutation does not affect IL-1α release in an in vivo model of peritoneal inflammation

As our *in vitro* data pointed to enhanced expression and release levels of mNLS-IL-1α after LPS treatment, we next performed an LPS intraperitoneal injection to mimic the entrance of bacterial-derived compounds into the host and induce an inflammatory response (21) (Figure 4A). We tested whether immune cells isolated from LPS-injected WT or mNLS mice still exhibited the nuclear or cytosolic localisation of pro-IL-1α, consistent with our *in vitro* experiments. We observed that peritoneal immune cells obtained from the peritoneal cavity of WT mice after LPS injection presented a predominantly nuclear localisation of IL-1α, while immune cells isolated from mNLS mice presented a mainly cytosolic IL-1α localisation (Figure 4B, D). IL-1β exhibited a cytosolic localisation in cells from both WT and mNLS mice (Figure 4C, E).

**Figure 4.**
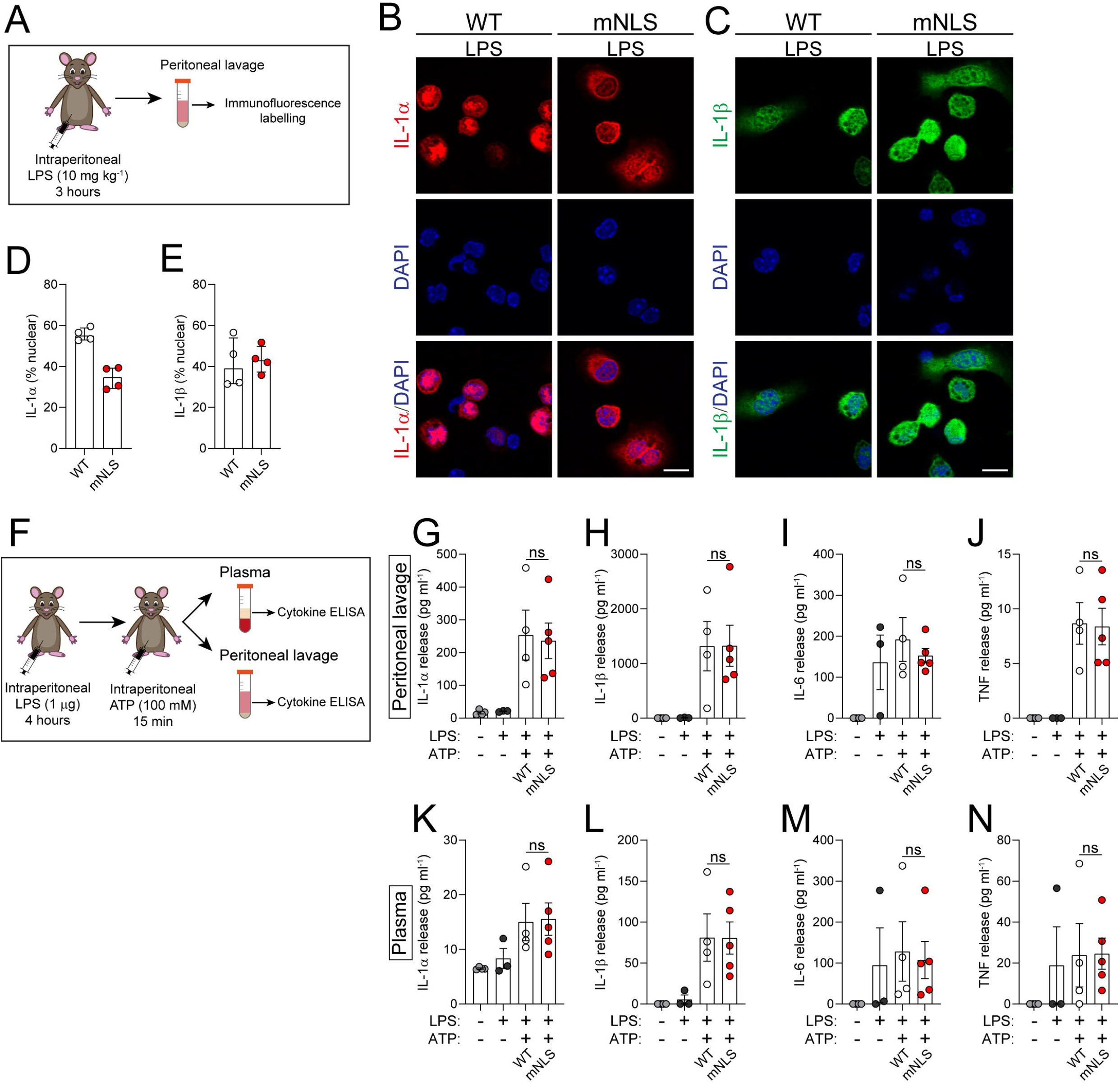
mNLS mutation did not affect IL-1α release in an *in vivo* model of peritoneal inflammation. **(A-E)** WT (n=2) and mNLS (n=2) mice were injected with LPS (10 mg/kg, intraperitoneal) for 3 hours, after which peritoneal lavage was collected. (A) Schematic of LPS injection model. Peritoneal lavage cells were analysed by immunofluorescence labelling for (B) pro-IL-1α and (C) pro-IL-1β, and nuclear localisation of (D) pro-IL-1α and (E) pro-IL-1β was quantified. A single z-plane is shown. Scale bar is 10 µm. **(F-N)** WT (n=4) and mNLS (n=5) mice were injected with LPS (1 µg, intraperitoneal) for 4 hours, followed by ATP (100 mM, intraperitoneal under anaethesia), after which plasma and peritoneal lavage were collected. Heterozygous mice were left untreated (n=4) or injected with LPS (1 µg, intraperitoneal) for 4 hours, followed by PBS (n=3). (F) Schematic of LPS+ATP injection model. Cytokine release was measured in (G-J) plasma and (K-N) peritoneal lavage. Data are presented as mean ± SEM. Data were analysed using unpaired t-test (G,H,I,J,L,N) or Mann-Whitney test (K,M). ns, not significant.

Next, we used an *in vivo* model of NLRP3-dependent inflammation where mice are administered with LPS followed by the NLRP3-activating stimulus ATP to induce NLRP3-dependent IL-1β release (22-24), and also IL-1α release, as we have previously reported when using this model (25). Thus, to determine whether mNLS mice secreted more IL-1α in response to inflammatory stimuli *in vivo* compared to WT mice, we injected mice i.p. with LPS followed by ATP and assayed the peritoneal lavage fluid and the plasma for IL-1α and other cytokines (Figure 4F). In the peritoneal lavage fluid, LPS and ATP increased the release of IL-1α and IL-1β from WT and mNLS mice, and there were no differences between the strains (Figure 4G, H). Likewise, induced levels of IL-6 and TNFα were the same across both strains (Figure 4I, J), and this was also reflected in the plasma (Figure 4K-N). Thus, under these conditions the release of IL-1α was not influenced by the NLS.

### Immune response to influenza infection in vivo shows no altered response in mNLS mice compared to WT

We had hypothesised that loss of IL-1α nuclear localisation might influence outcomes of viral infection. Here, we used a model of influenza infection, where it is known that IL-1α plays an important role in the modulation of the response to this pathogen (26). For this experiment, WT or mNLS mice were infected with influenza virus X31 for seven days, as we know at this timepoint there is an increase in viral load and reduced T cell numbers in IL-1R1 KO mice compared to WT (27) (Figure 5A). We also included uninfected heterozygote mice as controls. Influenza infection caused a gradual loss of weight over seven days, indicating progression of the infection, however infected WT and mNLS mice lost equivalent amounts of weight over this period (Figure 5B). When viral load was measured in the lung at day 7, no statistically significant differences were detected between WT and mNLS mice (Fig 5C). As expected, different immune cell subsets were recruited to the lung at seven days post-infection (Supplementary Figure 5 and 6), however, as with previous markers no significant differences were found between WT and mNLS infected mice.

**Figure 5.**
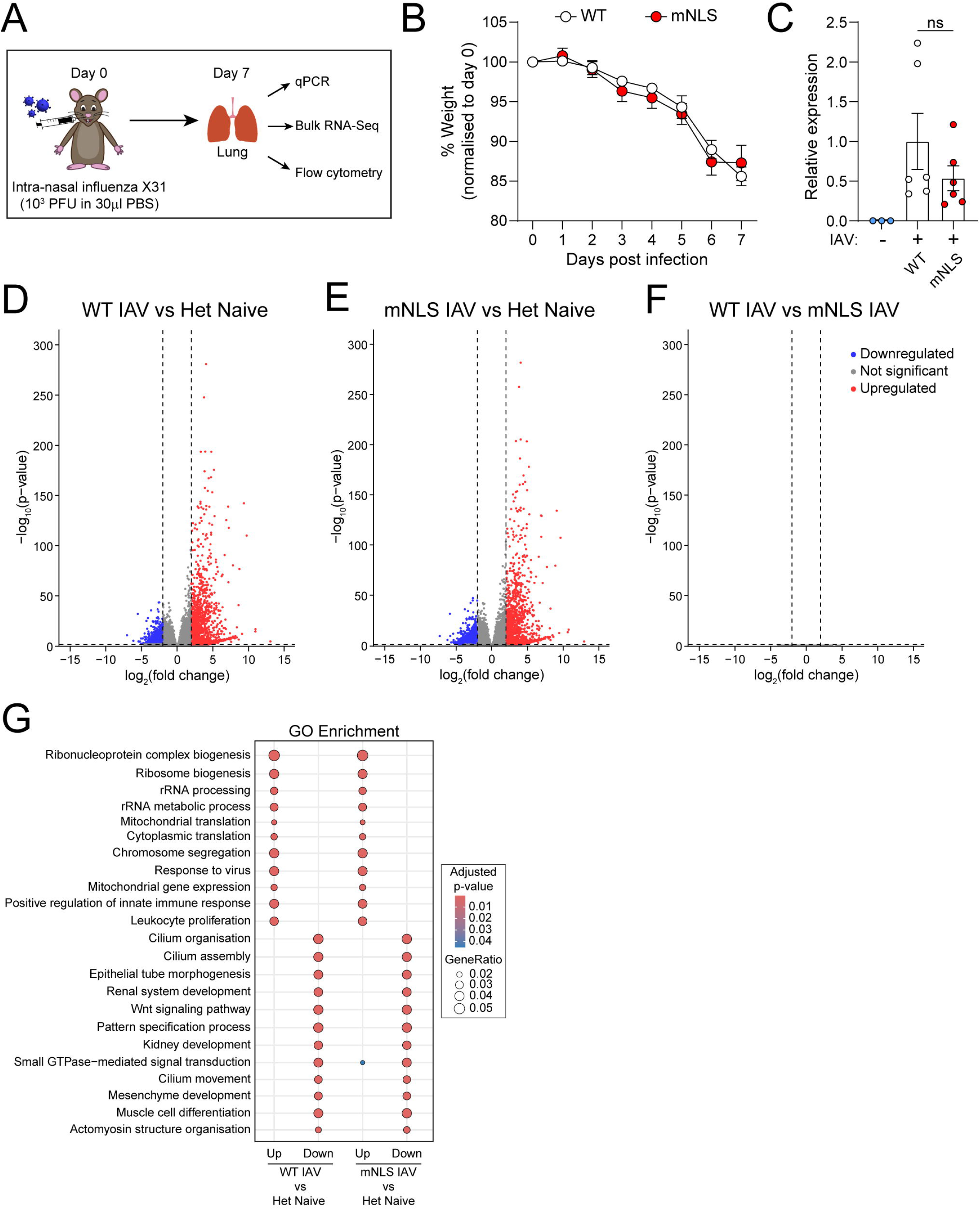
Pro-IL-1α mutation does not modulate the immune response in an influenza infection model. (A) On day 0, WT (n=6) or mNLS (n=6) mice were infected intranasally with live influenza strain X31 (10^3^ PFU in 30 µl of PBS), while heterozygous mice (n=3) were left uninfected (naïve) as a control. (B) Weight loss was monitored between days 0 and 7. (C) On day 7, the mice were culled, and viral load in the lungs was measured via qPCR. (D-F) Bulk RNA-Seq was performed on lung samples to assess gene expression, with differences between the groups expressed using volcano plots. The groups compared were (D) infected WT vs uninfected heterozygous, (E) infected mNLS vs uninfected heterozygous, and (F) infected WT vs infected mNLS. (G) Gene ontology (GO) enrichment analysis of biological processes for differentially expressed genes in WT IAV vs Het naïve (left) and mNLS IAV vs Het naïve (right).

As pro-IL-1α could have a transcriptional role via HAT activity regulation (3), and enhanced IL-1α secretion could impact the transcriptional profile via IL-1R1, we conducted an RNA-Seq analysis of lung tissue from uninfected heterozygote controls, and X31-infected WT and mNLS mice. The transcriptional profile observed in the lungs of infected WT and mNLS mice was strongly antiviral, as expected (Figure 5D, E and G), but no transcriptional differences where found between the strains. These data suggest that loss of IL-1α nuclear localisation does not provide an immune advantage in the protection from influenza infection, at least at the timepoint tested here.

## Discussion

IL-1α, unlike IL-1β, localises to the cell nucleus, driven by the presence of an NLS in its pro-domain. Here, we generated a mouse strain that contains two mutations in this NLS (mNLS), rendering its nuclear transport inactive. We validated this loss of nuclear localisation both *in vitro* and *ex vivo* and confirmed that upon LPS treatment, mNLS IL-1α was present in the cytosol while IL-1β localisation was not affected. Similarly, our data showed that IL-1α levels were not altered by the mNLS at the transcriptional level but were increased at the protein level, suggesting that NLS mutations might affect protein stability.

Nuclear IL-1α has been proposed to be a transcriptional regulator that can bind to HATs (3, 5) and is able to regulate inflammatory gene expression, including IL-6 (17). However, in our experimental models no changes in IL-6, TNF-α or any other transcriptional changes were observed after LPS priming in mNLS mice. One possibility is that any transcriptional changes mediated by pro-IL-1α in response to LPS may occur at later time points, as pro-IL-1α itself will first need to accumulate in the cell.

IL-1α plays an important role in response to pathogenic infections, including viruses. IL-1α is a key mediator of the innate immune response triggered by dsDNA in adenoviruses, and also in the response to IAV infection, by triggering induction of IL-1β and the consequent immune response to infection (26). We speculated that loss of the IL-1α NLS, as seen in species such as toothed whales, may modify the immune response to infection (2, 5). Toothed whales have lost functional mx1 and mx2 genes from their genome (28), which encode MxA and MxB respectively, two dynamin-like GTPase proteins that are important for anti-viral responses (29). The authors of this study hypothesised that the evolutionary pressure driving the loss of Mx proteins from the toothed whale arose from a virus that exploited Mx function, such as an ancestor of the herpes simplex virus 1 (28), and that in order to compensate for the loss of Mx proteins, other mutations may have occurred elsewhere in the toothed whale genome, which we speculated could be mutation of the NLS in pro-IL-1α. However, in our X31 influenza infection model we did not detect any significant differences in weight loss, or in the transcriptional profile in the lung, between WT and mNLS mice. Infected mNLS mice showed a trend towards expressing a higher amount of the T cell activation co-stimulatory markers CD80 and CD86 compared to infected WT mice in alveolar macrophages (Figure S5) and viral load showed a trend to be reduced in mNLS, however, these were not statistically significant. It is possible that effects of IL-1α in the mNLS mice could be more important at different time points post-infection. For instance, a reduction in IL-1β secretion levels in IL-1α KO mice was detected at 3 days post-infection (26) where there is more cellular damage, compared to our measurements that were obtained at 7 days post-infection. Similarly, it is possible that the nuclear role of IL-1α leads to epigenetic changes that might not impact the outcome of an initial infection, but that affect the response to a subsequent infection or insult.

One proposed explanation of why pro-IL-1α is localised to the nucleus is to prevent excessive release upon necrotic cell death, limiting the impact of the alarmin in the microenvironment (11). When we assessed release of IL-1α in response to both inflammasome-dependent and-independent activation stimuli, we observed a trend of increased release of IL-1α from mNLS BMDMs compared to WT. However, although no differences in transcriptional IL-1α levels were observed upon LPS priming, higher levels of pro-IL-1α protein levels were observed in mNLS BMDMs. This could explain the increase in IL-1α release from mNLS BMDMs described above.

Pro-IL-1α can be modified by different post-translational modifications (PTMs) including phosphorylation, ubiquitination and acetylation (30-32). Ubiquitination of pro-IL-1α contributes to protein stability, however the sites responsible for this modification have not been identified (31). Hence, it is possible that K86N mutation within the mNLS motif ^85^KNRW^88^ resulted in decreased ubiquitination of pro-IL-1α, impacting proteasomal degradation and hence affecting IL-1α stability, resulting in higher protein levels of pro-IL-1α in mNLS BMDMs. Acetylation has been described at K82 within the NLS and proposed to contribute to the nuclear localisation of the protein, since the mutation K82R (acetylation-null version) prevented its nuclear localisation and resulted in higher secretion levels (32). Interestingly, acetylation can also contribute to protein stability, and HATs such as p300/CBP can mediate the assembly of polyubiquitin chains (33), hence HAT binding to pro-IL-1α could also be an important regulator for protein stability in addition to histone modification and gene transcription.

Our work has mainly focused on the role of nuclear IL-1α in macrophages as these cells are important producers of these cytokines. However pro-IL-1α also plays important roles in non-immune cells such as keratinocytes, where secretion of IL-1α leads to an IFN driven signature on skin fibroblasts during viral infection with PRR-evasive viruses such as VSV (13). Hence other inflammatory models involving non-immune cells and IL-1α should be performed.

Overall, we have generated a new tool to investigate the nuclear role of IL-1α *in vitro* and *in vivo*. Although current experimental models tested here showed no apparent effect on the immune response in the absence of nuclear IL-1α, it is possible that nuclear localisation plays a more relevant role at earlier points in infection or in other inflammatory processes where more subtle roles of IL-1α are required, such as senescence or skin pathologies.

## Materials and methods

### Reagents

Goat anti-mouse IL-1α (AF-400-NA), goat anti-mouse IL-1β (AF-401-NA), mouse IL-1α ELISA (DY400), mouse IL-1β ELISA (DY401), mouse IL-6 ELISA (DY406) and mouse TNF ELISA (DY410) were from R&D Systems. LPS (in vitro: L2654; in vivo: L4516), ATP (A2383), nigericin (N7143), MCC950 (PZ0280), ZVAD (627610), necrostatin-1 (N9037), bovine serum albumin (A9647), (Ethylenedinitrilo)tetraacetic acid disodium salt (03690), Deoxyribonuclease I (D4513), Red Blood Cell Lysing Buffer Hybri-Max (R7757), formalin solution, neutral buffered, 10% (HT501128) and anti-β-actin-peroxidase (A3854) were from Sigma-Aldrich. Lipofectamine 3000, Alexa Fluor™ 594 donkey anti-goat IgG (A-11058), DAPI (D1306), ProLong Gold antifade mounting reagent (P36934), eBioscience™ Foxp3 / Transcription Factor Fixation/Permeabilization Concentrate and Diluent (00-5521-00), PureLink RNA mini kit (12183018A) and PureLink DNase set (12185010) were from Invitrogen. YVAD (4018838) was from Cambridge Bioscience. Superscript III Reverse Transcriptase (18080044), Power SYBR™ Green PCR Master Mix (4368706), CountBright™ Absolute Counting Beads (C36950) and RNAlater (AM7020) were from Thermo. Pam3CSK4 (tlrl-pms) and imiquimod (tlrl-imq) were from Invivogen. Cytiva Amersham ECL Prime Western Blotting Detection Reagent (RPN2236) was from GE Healthcare. Protease inhibitor cocktail (539131), Collagenase D (11088858001), DNase I (11284932001) and formaldehyde solution (252549) were from Merck. CytoTox 96 nonradioactive cytotoxicity assay (G1780) was from Promega. Rabbit anti-goat IgG (P044901-2) secondary antibody was from Agilent. Ionomycin calcium salt (CAY11932) was from Cayman Chemical. Collagenase, Type I (17100017) was from Gibco. RNeasy mini kit (74106) and HPRT1 QuantiTect primer assay (QT00059066) were from Qiagen. High-Capacity RNA-to-cDNA™ Kit (4387406) was from Applied Biosystems. BD Horizon™ Brilliant Stain Buffer Plus (566385), BD Pharmingen™ Purified Rat Anti-Mouse CD16/CD32 (553142, 2.4G2), BD Horizon™ PE-CF594 Hamster Anti-Mouse γδ T-Cell Receptor (563532, E50-2440), BD Horizon™ PE-CF594 Rat Anti-Mouse Siglec-F (562757, E50-2440), BD Horizon™ BUV395 Hamster Anti-Mouse TCR β Chain (569248, H57-597), BD Horizon™ BUV496 Rat Anti-Mouse CD4 (612952, GK1.5), BD Horizon™ BUV737 Rat Anti-CD11b (612800, M1/70) and BD Horizon™ BUV805 Rat Anti-Mouse CD45 (568336, 30-F11) were from BD Biosciences. Zombie UV™ Fixable Viability Kit (423108), Alexa Fluor® 488 anti-mouse CD80 (104716, 16-10A1), PerCP/Cyanine5.5 anti-mouse Ly-6C (128012, HK1.4), APC anti-mouse CD19 (115512, 6D5), Alexa Fluor® 700 anti-mouse Ly-6G (127622, 1A8), APC/Cyanine7 anti-mouse NK-1.1 (108724, PK136), Brilliant Violet 421™ anti-mouse CD64 (FcγRI) (164407, W18349C), Brilliant Violet 510™ anti-mouse/rat XCR1 (148218, ZET), Brilliant Violet 605™ anti-mouse CD86 (105037, GL-1), Brilliant Violet 650™ anti-mouse I-A/I-E (107641, M5/114.15.2), Brilliant Violet 711™ anti-mouse CD24 (101851, M1/69), Brilliant Violet 785™ anti-mouse CD11c (117336, N418), PE anti-mouse CD8a (100708, 53-6.7), PE/Cyanine7 anti-mouse F4/80 (123114, BM8) were from BioLegend.

### mNLS mice generation

Mice were maintained under standard laboratory conditions: ambient temperatures of 21°C (± 2°C), humidity of 40–50%, 12 h light cycle, ad libitum access to food and water. All procedures were performed blinded to genotype, and were carried out in accordance with the United Kingdom Animals (Scientific Procedures) Act 1986 and approved by the Home Office and the local Animal Ethical Review Group, University of Manchester.

sgRNA ggaagattctgaagaagaga which targets the NLS encoding residues of mIL1a was purchased as a chemically synthesised Alt-R oligo (IDT). An alt-R ssODN repair template (ttcaaggagagccgggtgacagtatcagcaacgtcaagcaacgggaagattctgaagaa*C*aga*T*ggctgagtttcagtgagaccttcact gaagatgacctgcagtccataacccatgat) comprising 60nt homology each side of the NLS sequences was designed to direct two base substitutions (italicised capitals) to mutate the motif KKRR to KNRW. Both sgRNA and ssODN were resuspended in sterile, RNase free injection buffer (TrisHCl 1mM, pH 7.5, EDTA 0.1mM) to a concentration of 100 ng/ml. 1 mg of sgRNA guide was mixed with 1 mg Cas9 recombinant protein (NEB) and incubated at RT for 10 mins, before ssODN was added to the injection mix (final concentrations 40 ng/ml; 40ng/ml respectively, ssODN HDR template (50 mg/ml)). Cryopreserved 1-cell C57BL/6J embryos from IVF were thawed and cultured for 4 hours, before they were pronuclear microinjected with the above mix using standard protocols. Zygotes were cultured overnight, and the resulting 2 cell embryos surgically implanted into the oviduct of day 0.5 post-coitum pseudopregnant mice. After birth and weaning genomic DNA was extracted from ear punches using Sigma REDExtract-N-Amp Tissue PCR kit and used to genotype pups by amplifying across the NLS sequence with primers DB27 Geno F1 tcaggttccacttttcctcct and DB27 Geno R1 tctttccactccattctaccact. Amplicons were Sanger sequenced to identify mice harbouring the mutated allele, and a founder mouse confirmed and colony established.

### In vivo peritoneal inflammation models

For the LPS-induced peritoneal inflammation model, WT or mNLS mice were administered with LPS (10 mg/kg, from Escherichia coli 0127:B8) intraperitoneally (21). Three hours after injection, mice were anesthetised with isoflurane, and the peritoneal cavity was lavaged with 3 ml of RPMI media. Peritoneal lavage was centrifuged at 1500 x g for 5 min at 4°C, and the cell pellet was resuspended in DMEM and seeded onto coverslips, left to adhere for 1.5 h, and then fixed and assessed for IL-1α and IL-1β subcellular location by immunofluorescence labelling.

For the LPS and ATP-induced peritoneal inflammation model, WT or mNLS mice were administered with LPS (1 ug, from Escherichia coli 0127:B8) intraperitoneally (25). Four hours after injection, mice were anesthetised with isoflurane, and injected with ATP (100 mM, pH 7.4) for 15 min. Peritoneal lavage was collected as above, and blood was collected via cardiac puncture. Peritoneal lavage was centrifuged at 1500 x g for 5 min at 4°C and the supernatant was analysed for cytokine content. Blood was centrifuged at 1500 x g for 15 min at 4°C, and the supernatant was subsequently centrifuged at 18000 x g for 3 min at 4°C. The resulting supernatant (plasma) was assessed for cytokine content.

### In vivo influenza infection model

WT or mNLS mice were anaesthetized by 2.5% inhalation isoflurane and intranasally infected with 10^3^ PFU of the IAV strain X31 (H3N2), in a volume of 30 µL PBS, on day 0. Mild to moderate weight loss were expected from IAV X31 infection, and daily weights were monitored until day 7 post-infection. Infected WT and mNLS mice, and heterozygous control mice, were culled on day 7 post-IAV X31 infection. Lung draining (cervical and mediastinal) lymph nodes were first collected, and after PBS perfusion with 10 mL of 1X PBS 1 mM EDTA, three lobes of lung and spleen were collected. The lung and spleen were weighed, minced, and digested with either collagenase type I (10 mg/mL) and DNase I (50 µg/mL) for 40 minutes (lungs), or with collagenase type D (1 mg/uL) and DNase I (50 µg/mL) for 30 minutes (spleens), at 37°C in a shaking incubator. The lung draining lymph nodes were counted and sized. The digested lung and spleen tissue, and lung draining lymph nodes, were passed through a 70 µm filter, washed with 25 mL of cold PBS 5 mM EDTA solution, and centrifuged at 500 xg for 5 minutes. Pelleted cells were resuspended in 1 mL of RBC lysis buffer and washed with 10 mL of PBS. The RBC-lysed single cell suspension was resuspended in complete RPMI media with 10% FCS before flow cytometry staining.

### Flow Cytometry

Single cell suspensions (1/5^th^ of lung) were transferred to 5 mL round-bottom polystyrene (FACS) tubes, washed with 3 mL 1x PBS, and resuspended in 300 µL of Zombie UV Live/Dead stain. After 15 minutes, cells were washed with 3 mL 1x PBS and resuspended in 50 µL of anti-CD16/32 (Fc block) in FACS buffer (2% FCS in PBS 2 mM EDTA). After 10 minutes, without washing, 10 µL of antibody staining mastermix prepared in BD Brilliant Stain Buffer Plus was added to samples, and incubated for 30 minutes at 4°C in the dark. Stained cells were washed with 3 mL 1x PBS, and cells were fixed with eBioscience™ Foxp3 / Transcription Factor Staining Buffer Kit as per manufacturer’s guidelines. The fixed samples were passed through a 35 µm filter and resuspended in 300 µL of 1x PBS. 25 µL of counting beads were spiked into each sample before flow cytometry acquisition, to calculate cell numbers per 1 g of tissue. Gating strategy is shown in Supplementary Figure 5.

### Cell culture

Primary bone marrow-derived macrophages (BMDMs) were prepared from the femurs and tibias of 3-6-month-old male and female WT and mNLS C57BL/6 mice. Bone marrow was isolated, red blood cells were lysed, and the remaining cells were passed through a cell strainer Cells were cultured in 70% DMEM (containing 10% FBS, penicillin (100 U ml^-1^) and streptomycin (100 ug ml^-1^)) supplemented with L929-conditioned medium for 7 days at 37°C and 5% CO2. Cells were seeded at a density of 1 x 10^6^ cells ml^-1^ for experiments and left to adhere overnight.

Peritoneal macrophages were isolated from the peritoneal lavage. Mice were culled by cervical dislocation, and the peritoneal cavity was lavaged with 5 ml RPMI (containing 3% FBS, 1 mM EDTA). The collected RPMI was centrifuged at 1500 x g, 5 min, and the cell pellet resuspended in fresh DMEM (containing 10% FBS, penicillin (100 U ml^-1^) and streptomycin (100 ug ml^-1^)). Cells were seeded at a density of 1 x 10^6^ cells ml^-1^ for experiments and left to adhere overnight.

### Cell treatment protocols

For IL-1α priming experiments, WT or mNLS BMDMs or peritoneal macrophages were treated with vehicle (PBS) or LPS (1 µg/ml) for 4 hours. Cell lysates and supernatants were subsequently analysed using ELISA, immunofluorescence labelling, qPCR and RNA-Seq.

For IL-1α processing and release experiments, WT or mNLS BMDMs were first primed with LPS (1 µg/ml) or Pam3CSK4 (100 ng/ml) for 4 hours. IL-1α processing was then induced by treatment with ionomycin (10 µM, 1 h), nigericin (10 μM, 1 h), ATP (5 mM, 1 h), transfection of LPS (2 μg/ml, 18 h) using lipofectamine 3000, or ZVAD (50 μM, 5 h) in the presence or absence of MCC950 (10 µM; MCC), calpeptin (40 µM; Calp), YVAD (100 μM), or necrostatin-1 (50 μM; Nec1). Cell lysates and supernatants were subsequently analysed using ELISA, western blotting, and cell death assays.

### RNA-Sequencing

BMDMs were primed with either vehicle (PBS) or LPS (1 µg ml^-1^) for 4 hours. Total RNA was extracted using a PureLink RNA mini kit. Total RNA was submitted to the Genomic Technologies Core Facility (GTCF) at the University of Manchester. Quality and integrity of the RNA samples were assessed using a 4200 TapeStation (Agilent Technologies) and then libraries generated using the *Illumina® Stranded mRNA Prep. Ligation* kit (Illumina, Inc.) according to the manufacturer’s protocol. Briefly, total RNA (typically 0.025-1ug) was used as input material from which polyadenylated mRNA was purified using poly-T, oligo-attached, magnetic beads. Next, the mRNA was fragmented under elevated temperature and then reverse transcribed into first strand cDNA using random hexamer primers and in the presence of Actinomycin D (thus improving strand specificity whilst mitigating spurious DNA-dependent synthesis). Following removal of the template RNA, second strand cDNA was then synthesised to yield blunt-ended, double-stranded cDNA fragments. Strand specificity was maintained by the incorporation of deoxyuridine triphosphate (dUTP) in place of dTTP to quench the second strand during subsequent amplification. Following a single adenine (A) base addition, adapters with a corresponding, complementary thymine (T) overhang were ligated to the cDNA fragments. Pre-index anchors were then ligated to the ends of the double-stranded cDNA fragments to prepare them for dual indexing. A subsequent PCR amplification step was then used to add the index adapter sequences to create the final cDNA library. The adapter indices enabled the multiplexing of the libraries, which were pooled prior to loading on to the appropriate flow-cell. This was then paired-end sequenced (59 + 59 cycles, plus indices) on an Illumina NovaSeq6000 instrument. Finally, the output data was demultiplexed and BCL-to-Fastq conversion performed using Illumina’s bcl2fastq software, version 2.20.0.422.

### Bioinformatic data analysis and visualisation

All analyses were conducted in R version 2025.05.1+513 (36). For principal component analysis (PCA), count matrices were imported into R and annotated with sample metadata (e.g., genotype and treatment groups). Genes lacking annotation were excluded. A variance stabilising transformation (VST) was applied using the DESeq2 package (37) to normalise raw counts and stabilise variance across expression levels. Principal component analysis was performed with the plotPCA function in DESeq2, and the first two principal components were visualised with ggplot2 (38), with the percentage of variance explained displayed on each axis. For volcano plots, differential expression results were imported from processed RNA-Seq datasets and pre-formatted to include gene identifiers, log_2_ fold changes, and adjusted *p*-values. Genes with missing information were excluded, and *p*-values equal to zero were replaced by the smallest positive floating-point value in R for numerical stability. Genes were classified as *upregulated* or *downregulated* using thresholds of |log_2_ fold change| ≥ 2 and *FDR*-adjusted *p*-value ≤ 0.05; all other genes were classified as not significant. Volcano plots were generated with ggplot2 and ggrepel (39), plotting log_2_ fold changes against – log_10_(*p*-adjusted values). Horizontal and vertical dotted lines indicated statistical significance and fold-change cutoffs, respectively. For gene ontology (GO) enrichment, for each dataset, significantly upregulated and downregulated genes (*FDR* ≤ 0.05) were converted from gene symbols to Entrez Gene identifiers using the org.Mm.eg.db annotation package (40). Enrichment analyses were conducted with the compareCluster function from clusterProfiler (41), specifying the biological process (BP) ontology. Significant enrichment was defined at *p* ≤ 0.05. Results were visualised as dot plots, highlighting the most enriched terms.

### qPCR

BMDMs from WT and mNLS mice were treated as described above and RNA extracted as done for the RNA-Seq experiment. RNA (300 ng) was converted to cDNA using SuperScript III reverse transcriptase according to manufacturer’s instructions. qPCR was performed using Power SYBR Green PCR master mix and 200 nM of each primer using a 7900HT Fast Real-Time PCR System (Applied Biosystems), and 3 μl of 1:10-diluted cDNA was loaded. Samples were run in triplicate. Data were normalised to the expression of the housekeeping gene *Hmbs*. Expression levels of genes of interest were computed as follows: relative mRNA expression = *E*^−(*C*t of gene of interest)^/ *E*^−(*C*t of housekeeping gene)^, where *C*t is the threshold cycle value and *E* is efficiency. The following primers were used: IL-1α forward, TCTCAGATTCACAACTGTTCGTG, IL-1α reverse, AGAAAATGAGGTCGGTCTCACTA; IL-1β forward, AACCTGCTGGTGTGTGACGTTC, IL-1β reverse, CAGCACGAGGCTTTTTTGTTGT; Hmbs forward, GAAATCATTGCTATGTCCACCA, Hmbs reverse, GCGTTTTCTAGCTCCTTGGTAA.

To determine lung viral load, one lung lobe from WT and mNLS mice infected with IAV, and uninfected heterozygous mice, was collected in a sterile 1.5 mL tube containing RNALater and was later transferred into a 2 mL tube containing RLT buffer supplemented with β-mercaptoethanol and a single 2 mm stainless steel bead. Tissue was homogenised into suspension using a Qiagen Tissue Lyser. Total RNA was then extracted using the RNeasy Mini Kit according to the manufacturer’s protocol. Subsequently, the purified RNA was reverse-transcribed into cDNA using the High-Capacity RNA-to-cDNA™ Kit, following the manufacturer’s instructions. Viral cDNA was quantified using a TaqMan real-time PCR assay on a QuantStudio™ 12K Flex Real-Time PCR System (Applied Biosystems). The assay was designed to quantify the influenza A matrix protein (MP) gene using the following primers and probe: MP forward, AAGACCAATCCTGTCACCTCTGA, MP reverse, CAAAGCGTCTACGCTGCAGTCC, MP probe: [6-FAM]-TTTGTGTTCACGCTCACCGTT-[TAMRA]. All samples were analysed in triplicate. Gene expression levels were normalised to the endogenous control gene HPRT1. Relative gene expression was calculated using the comparative threshold cycle (Ct) method. The fold change in target gene expression was determined using the formula 2−^ΔΔ Ct^, which represents the expression level relative to that of untreated control samples.

### Cell death assay

Cell death was determined by measuring lactate dehydrogenase release into the supernatant using a CytoTox 96 Non-Radioactive Cytotoxicity assay (Promega), according to the manufacturer’s instructions.

### ELISA

The levels of IL-1α, IL-1β, IL-6 and TNF in the supernatant were measured by ELISA according to the manufacturer’s instructions.

### Immunofluorescence assays

Cells were washed once in PBS then fixed in 4% paraformaldehyde (PFA) for 10 min before being incubated in blocking solution (5% BSA, PBS, 0.1% Triton X-100) for 1 h at room temperature. Cells were incubated with goat anti-mouse IL-1α (2 μg ml^-1^) or goat anti-mouse IL-1β (2 μg ml^-1^) primary antibodies in 5% BSA, PBS, 0.1% Triton X-100 at 4 °C overnight. Cells were then washed and incubated with Alexa Fluor™ 594 donkey anti-goat IgG (4 µg ml^-1^) secondary antibody for 1 h at room temperature. Nuclei were stained with DAPI (0.5 µg ml^-1^). Coverslips were mounted on slides using ProLong Gold antifade mounting reagent and left to dry overnight.

### Confocal microscopy

Confocal microscopy images were acquired using a 63×/1.40 HCS PL Apo objective on a Leica TCS SP8 AOBS upright confocal microscope with LAS X software (v3.5.1.18803). To prevent interference between channels, lasers were excited sequentially for each channel. The blue diode with 405 nm and the white light laser with the 594 nm laser line was used, with hybrid and photon-multiplying tube detectors with detection mirror settings set appropriately. Z-stacks were acquired with 0.3 µm steps between Z sections. Images were acquired from 3-5 fields of view from each independent experiment.

### Image analysis

Nuclear localisation was quantified manually on FIJI (ImageJ) using maximum projections and expressed as the percentage of total pro-IL-1α or pro-IL-1β fluorescence intensity that co-localised with DAPI signal.

### Western blotting

Supernatant was collected and cells were lysed in lysis buffer (50 mM Tris-HCl, 150_JmM NaCl; Triton-X-100 1% v/v, pH 7.3) containing protease inhibitor cocktail. Lysates were analysed for IL-1α and IL-1β. Equal volumes of lysates or supernatants were loaded. Samples were run on SDS-polyacrylamide gels and transferred at 25 V onto nitrocellulose or PVDF membranes using a Trans-Blot® Turbo Transfer™ System (Bio-Rad). Membranes were blocked in 5% BSA (w/v) or 5% milk (w/v) in PBS, 0.1% Tween (v/v) (PBST) for 1 h at room temperature. Membranes were incubated at 4°C overnight with goat anti-mouse IL-1α (200 ng/ml) or goat anti-mouse IL-1β (250 ng/ml) primary antibodies. Membranes were washed in PBST and incubated with rabbit anti-goat IgG (500 ng/ml) in 5% BSA (w/v) in PBST at room temperature for 1 h. Proteins were visualised with Cytiva Amersham ECL Prime Western Blotting Detection Reagent and G:BOX (Syngene) and Genesys software. Membranes were probed for β-actin as a loading control. Densitometry performed on cell lysates was first normalised to β-actin, and then expressed relative to a control treatment. Densitometry performed on supernatants was not normalised.

### Data analysis

Nuclear localisation quantification from immunofluorescence images is presented as median ± interquartile range, with each data point representing a field of view. All other data are presented as mean ± SEM. Data were assessed for normal distribution using Shapiro-Wilk normality test. Parametric data were analysed using unpaired t test or one-way ANOVA with Dunnett’s or Sidak’s post-hoc test. Non-parametric data were analysed using unpaired two-tailed Mann-Whitney test or Kruskal-Wallis test with Dunn’s post-hoc test. Western blots are representative of 6-7 independent experiments. Statistical analysis was performed using GraphPad Prism (v10).

## Data availability statement

Data is available from the authors upon reasonable request.

## Acknowledgements

BBSRC project grant (BB/Y004876/1) (GLC, DB, CH, RD-P). The Dowager Countess Eleanor Peel Trust Sir Robert Boyd fellowship (CH). Medical Research Council (MRC) programme grant (DB and CH, MR/T016515/1).

We thank Leo Zeef, Andy Hayes and Michal Smiga of the Bioinformatics and Genomic Technologies Core Facilities at the University of Manchester for providing support with RNA-Sequencing, Dr. Gareth Howell and the University of Manchester Flow Cytometry Core Facility for facilitating flow cytometry analysis, and members of the Biological Service Facility at the University of Manchester for help with animal work. The Bioimaging Facility microscopes used in this study were purchased with grants from BBSRC, Wellcome, and the University of Manchester Strategic Fund.

## Author contributions

Conceptualisation: CH, DB, GL-C

Methodology: MT, AA, GC, CL

Investigation: CH, RDP, SML, JPG

Writing – original draft: CH, DB, GL-C

Writing – review and editing: CH, RDP, DB, GL-C, MT

Visualisation: CH, RDP, SML

Funding acquisition: CH, DB, GL-C

Supervision: DB, GL-C

Resources: CBL

## References

1. Cavalli G, Colafrancesco S, Emmi G, Imazio M, Lopalco G, Maggio MC, et al. Interleukin 1α: a comprehensive review on the role of IL-1α in the pathogenesis and treatment of autoimmune and inflammatory diseases. Autoimmunity reviews. 2021;20(3):102763.

2. Rivers-Auty J, Daniels MJD, Colliver I, Robertson DL, Brough D. Redefining the ancestral origins of the interleukin-1 superfamily. Nat Commun. 2018;9(1):1156.

3. Buryskova M, Pospisek M, Grothey A, Simmet T, Burysek L. Intracellular interleukin-1alpha functionally interacts with histone acetyltransferase complexes. J Biol Chem. 2004;279(6):4017–26.

4. Zamostna B, Novak J, Vopalensky V, Masek T, Burysek L, Pospisek M. N-terminal domain of nuclear IL-1α shows structural similarity to the C-terminal domain of Snf1 and binds to the HAT/core module of the SAGA complex. PLoS One. 2012;7(8):e41801.

5. Wellens R, Tapia VS, Seoane PI, Bennett H, Adamson A, Coutts G, et al. Proximity labelling of pro-interleukin-1α reveals evolutionary conserved nuclear interactions. Nat Commun. 2024;15(1):6750.

6. Luheshi NM, Rothwell NJ, Brough D. The dynamics and mechanisms of interleukin-1alpha and beta nuclear import. Traffic (Copenhagen, Denmark). 2009;10(1):16–25.

7. Yamada A, Wake K, Imaoka S, Motoyoshi M, Yamamoto T, Asano M. Analysis of the effects of importin α1 on the nuclear translocation of IL-1α in HeLa cells. Sci Rep. 2024;14(1):1322.

8. Tapia VS, Daniels MJD, Palazón-Riquelme P, Dewhurst M, Luheshi NM, Rivers-Auty J, et al. The three cytokines IL-1β, IL-18, and IL-1α share related but distinct secretory routes. J Biol Chem. 2019;294(21):8325–35.

9. Wiggins KA, Parry AJ, Cassidy LD, Humphry M, Webster SJ, Goodall JC, et al. IL-1α cleavage by inflammatory caspases of the noncanonical inflammasome controls the senescence-associated secretory phenotype. Aging cell. 2019;18(3):e12946.

10. Afonina IS, Müller C, Martin SJ, Beyaert R. Proteolytic Processing of Interleukin-1 Family Cytokines: Variations on a Common Theme. Immunity. 2015;42(6):991–1004.

11. Luheshi NM, McColl BW, Brough D. Nuclear retention of IL-1 alpha by necrotic cells: a mechanism to dampen sterile inflammation. Eur J Immunol. 2009;39(11):2973–80.

12. Milora KA, Miller SL, Sanmiguel JC, Jensen LE. Interleukin-1α released from HSV-1-infected keratinocytes acts as a functional alarmin in the skin. Nat Commun. 2014;5:5230.

13. Orzalli MH, Smith A, Jurado KA, Iwasaki A, Garlick JA, Kagan JC. An Antiviral Branch of the IL-1 Signaling Pathway Restricts Immune-Evasive Virus Replication. Mol Cell. 2018;71(5):825-40.e6.

14. Orzalli MH, Prochera A, Payne L, Smith A, Garlick JA, Kagan JC. Virus-mediated inactivation of anti-apoptotic Bcl-2 family members promotes Gasdermin-E-dependent pyroptosis in barrier epithelial cells. Immunity. 2021;54(7):1447-62.e5.

15. Daniels MJD, Adamson AD, Humphreys N, Brough D. CRISPR/Cas9 mediated mutation of mouse IL-1α nuclear localisation sequence abolishes expression. Sci Rep. 2017;7(1):17077.

16. Chan J, Atianand M, Jiang Z, Carpenter S, Aiello D, Elling R, et al. Cutting Edge: A Natural Antisense Transcript, AS-IL1α, Controls Inducible Transcription of the Proinflammatory Cytokine IL-1α. Journal of immunology (Baltimore, Md: 1950). 2015;195(4):1359–63.

17. Werman A, Werman-Venkert R, White R, Lee JK, Werman B, Krelin Y, et al. The precursor form of IL-1alpha is an intracrine proinflammatory activator of transcription. Proc Natl Acad Sci U S A. 2004;101(8):2434–9.

18. Gross O, Yazdi AS, Thomas CJ, Masin M, Heinz LX, Guarda G, et al. Inflammasome activators induce interleukin-1α secretion via distinct pathways with differential requirement for the protease function of caspase-1. Immunity. 2012;36(3):388–400.

19. England H, Summersgill HR, Edye ME, Rothwell NJ, Brough D. Release of interleukin-1α or interleukin-1β depends on mechanism of cell death. J Biol Chem. 2014;289(23):15942–50.

20. Tsuchiya K, Hosojima S, Hara H, Kushiyama H, Mahib MR, Kinoshita T, Suda T. Gasdermin D mediates the maturation and release of IL-1α downstream of inflammasomes. Cell Rep. 2021;34(12):108887.

21. Baldwin AG, Rivers-Auty J, Daniels MJD, White CS, Schwalbe CH, Schilling T, et al. Boron-Based Inhibitors of the NLRP3 Inflammasome. Cell Chem Biol. 2017;24(11):1321-35.e5.

22. Griffiths RJ, Stam EJ, Downs JT, Otterness IG. ATP induces the release of IL-1 from LPS-primed cells in vivo. Journal of immunology (Baltimore, Md: 1950). 1995;154(6):2821–8.

23. Solle M, Labasi J, Perregaux DG, Stam E, Petrushova N, Koller BH, et al. Altered cytokine production in mice lacking P2X(7) receptors. J Biol Chem. 2001;276(1):125–32.

24. Perregaux DG, McNiff P, Laliberte R, Hawryluk N, Peurano H, Stam E, et al. Identification and characterization of a novel class of interleukin-1 post-translational processing inhibitors. J Pharmacol Exp Ther. 2001;299(1):187–97.

25. Daniels MJ, Rivers-Auty J, Schilling T, Spencer NG, Watremez W, Fasolino V, et al. Fenamate NSAIDs inhibit the NLRP3 inflammasome and protect against Alzheimer’s disease in rodent models. Nat Commun. 2016;7:12504.

26. Momota M, Lelliott P, Kubo A, Kusakabe T, Kobiyama K, Kuroda E, et al. ZBP1 governs the inflammasome-independent IL-1α and neutrophil inflammation that play a dual role in anti-influenza virus immunity. Int Immunol. 2020;32(3):203–12.

27. Schmitz N, Kurrer M, Bachmann MF, Kopf M. Interleukin-1 is responsible for acute lung immunopathology but increases survival of respiratory influenza virus infection. J Virol. 2005;79(10):6441–8.

28. Braun BA, Marcovitz A, Camp JG, Jia R, Bejerano G. Mx1 and Mx2 key antiviral proteins are surprisingly lost in toothed whales. Proc Natl Acad Sci U S A. 2015;112(26):8036–40.

29. Zav’yalov VP, Hämäläinen-Laanaya H, Korpela TK, Wahlroos T. Interferon-Inducible Myxovirus Resistance Proteins: Potential Biomarkers for Differentiating Viral from Bacterial Infections. Clin Chem. 2019;65(6):739–50.

30. Beuscher HU, Nickells MW, Colten HR. The precursor of interleukin-1 alpha is phosphorylated at residue serine 90. J Biol Chem. 1988;263(8):4023–8.

31. Ainscough JS, Frank Gerberick G, Zahedi-Nejad M, Lopez-Castejon G, Brough D, Kimber I, Dearman RJ. Dendritic cell IL-1α and IL-1β are polyubiquitinated and degraded by the proteasome. J Biol Chem. 2014;289(51):35582–92.

32. Cohen I, Rider P, Vornov E, Tomas M, Tudor C, Wegner M, et al. IL-1α is a DNA damage sensor linking genotoxic stress signaling to sterile inflammation and innate immunity. Sci Rep. 2015;5:14756.

33. Sadoul K, Boyault C, Pabion M, Khochbin S. Regulation of protein turnover by acetyltransferases and deacetylases. Biochimie. 2008;90(2):306–12.

34. Dinarello CA. Overview of the IL-1 family in innate inflammation and acquired immunity. Immunol Rev. 2018;281(1):8–27.

35. Cayrol C, Girard JP. Interleukin-33 (IL-33): A nuclear cytokine from the IL-1 family. Immunol Rev. 2018;281(1):154–68.

36. R Core Team. R: A Language and Environment for Statistical Computing. Vienna, Austria: R Foundation for Statistical Computing; 2025.

37. Love MI, Huber W, Anders S. Moderated estimation of fold change and dispersion for RNA-seq data with DESeq2. Genome Biol. 2014;15(12):550.

38. Wickham H. ggplot2: Elegant Graphics for Data Analysis. New York: Springer-Verlag; 2016.

39. Slowikowski K. ggrepel: Automatically Position Non-Overlapping Text Labels with ‘ggplot2’. The Comprehensive R Archive Network. 2024.

40. Carlson M. org.Mm.eg.db: Genome wide annotation for Mouse. R package. 2019.

41. Yu G, Wang LG, Han Y, He QY. clusterProfiler: an R package for comparing biological themes among gene clusters. Omics. 2012;16(5):284–7.

