## Supplementary Figures for "Inflammatory responses following CRISPR modification of the nuclear localisation sequence in endogenous interleukin-1α"

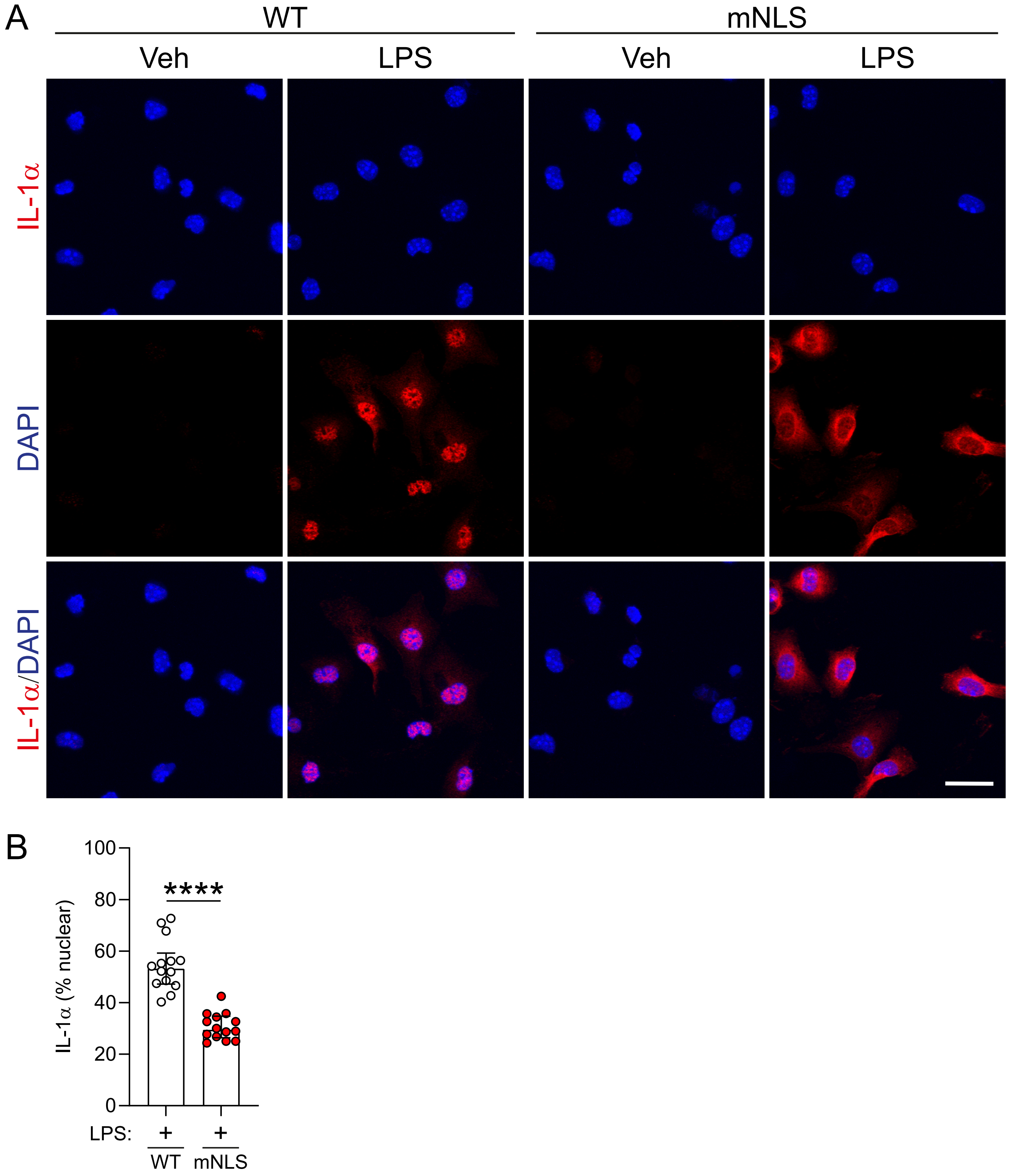


**Supplementary Figure 1. Pro-IL-1α NLS mutation reduces nuclear localisation in peritoneal macrophages.** Peritoneal macrophages were isolated from WT or mNLS mice, and primed with vehicle (PBS) or LPS (1 μg/ml, 4 h). (A) Immunofluorescence labelling of pro-IL-1α, and (B) quantification of nuclear localisation of pro-IL-1α (n=3 biological replicates, with 4-5 random fields of view per replicate). Scale bar is 20 μm. Data are presented as median ± IQR. Data were analysed using unpaired t-test. ****P<0.0001.

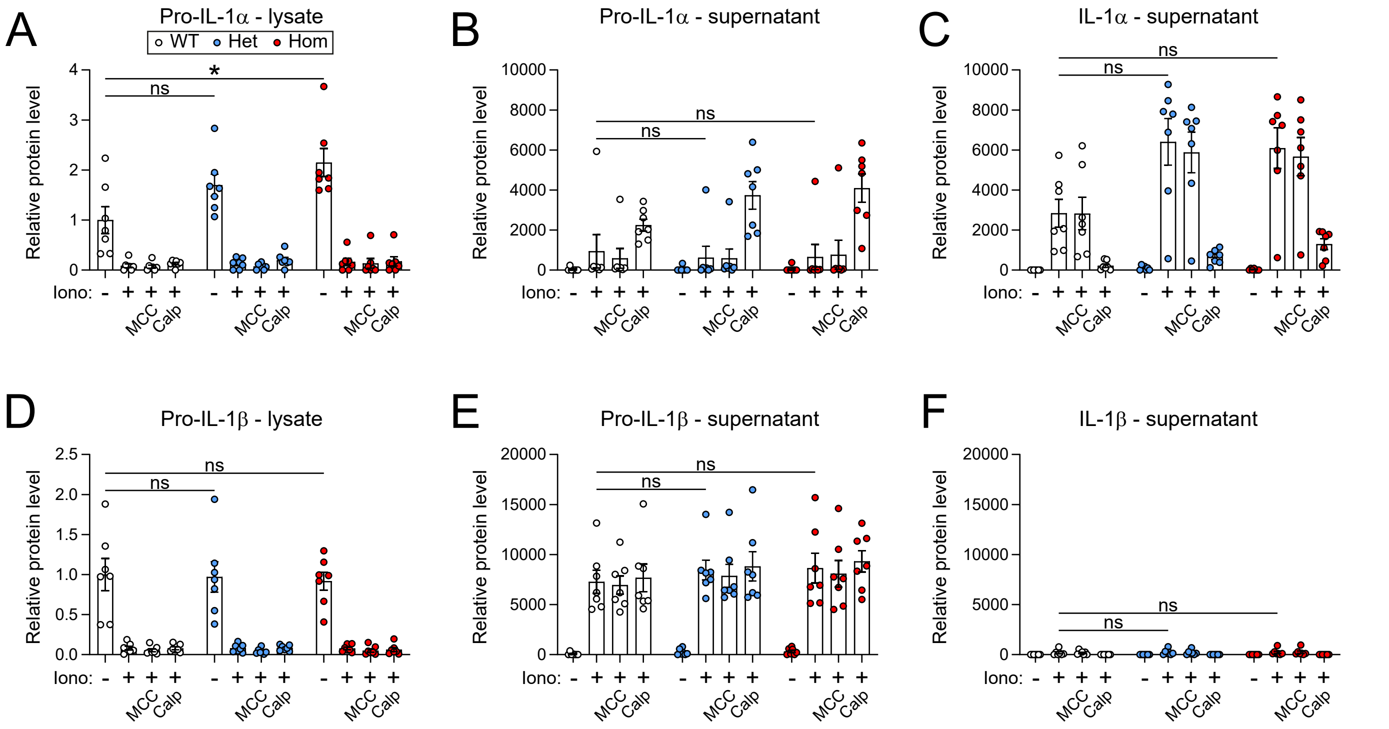


**Supplementary Figure 2. Pro-IL-1α NLS mutation does not negatively affect processing and release of IL-1α in response to ionomycin. (A-F)** Densitometry of western blots from Figure 2D (n=7). (A) Relative pro-IL-1α levels were determined in the lysate, and (B) pro-IL-1α and (C) mature IL-1α levels were determined in the supernatant. (D) Relative pro-IL-1β levels were determined in the lysate, and (E) pro-IL-1β and (F) mature IL-1β levels were determined in the supernatant. Data are presented as mean ± SEM. Data were analysed using Kruskal-Wallis test followed by Dunn's post-hoc test (A,B,C,E,F) or one-way ANOVA followed by Dunnett's post-hoc test (D). *$P<0.05$; ns, not significant.

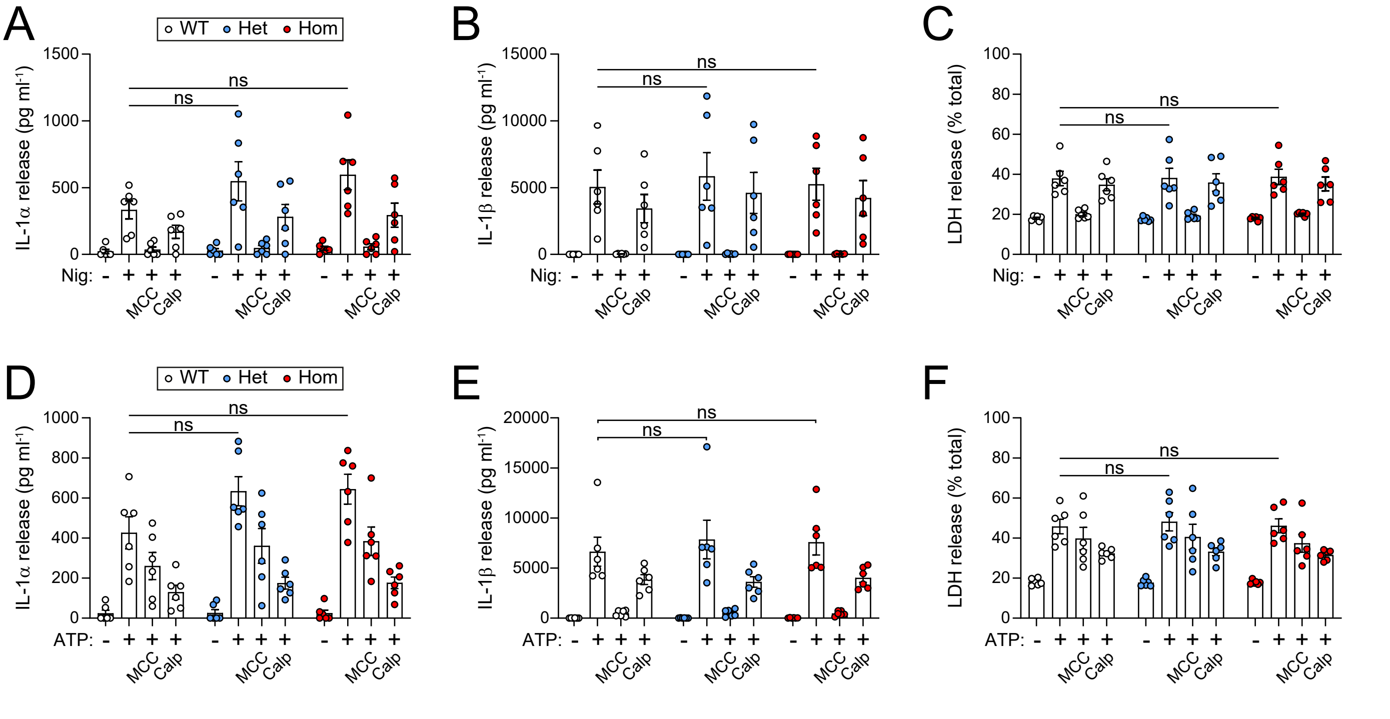


**Supplementary Figure 3. Pro-IL-1α NLS mutation does not negatively affect processing and release of IL-1α in response to ATP and nigericin. (A-C)** WT or mNLS BMDMs were primed with LPS (1 μg/ml, 4 h), followed by nigercin treatment (10 μM, 1 h) in the presence or absence of MCC950 (10 μM; MCC) or calpeptin (40 μM; Calp) (n=6). Supernatants were assessed for (A) IL-1α release, (B) IL-1β release and (C) LDH release. **(D-F)** WT or mNLS BMDMs were primed with LPS (1 μg/ml, 4 h), followed by ATP treatment (5 mM, 1 h) in the presence or absence of MCC950 (10 μM) or calpeptin (40 μM) (n=6). Supernatants were assessed for (D) IL-1α release, (E) IL-1β release and (F) LDH release. Data are presented as mean ± SEM. Data were analysed using one-way ANOVA followed by Dunnett's post-hoc test (A,B,D,F) or Kruskal-Wallis test followed by Dunn's post-hoc test (C,E). ns, not significant.

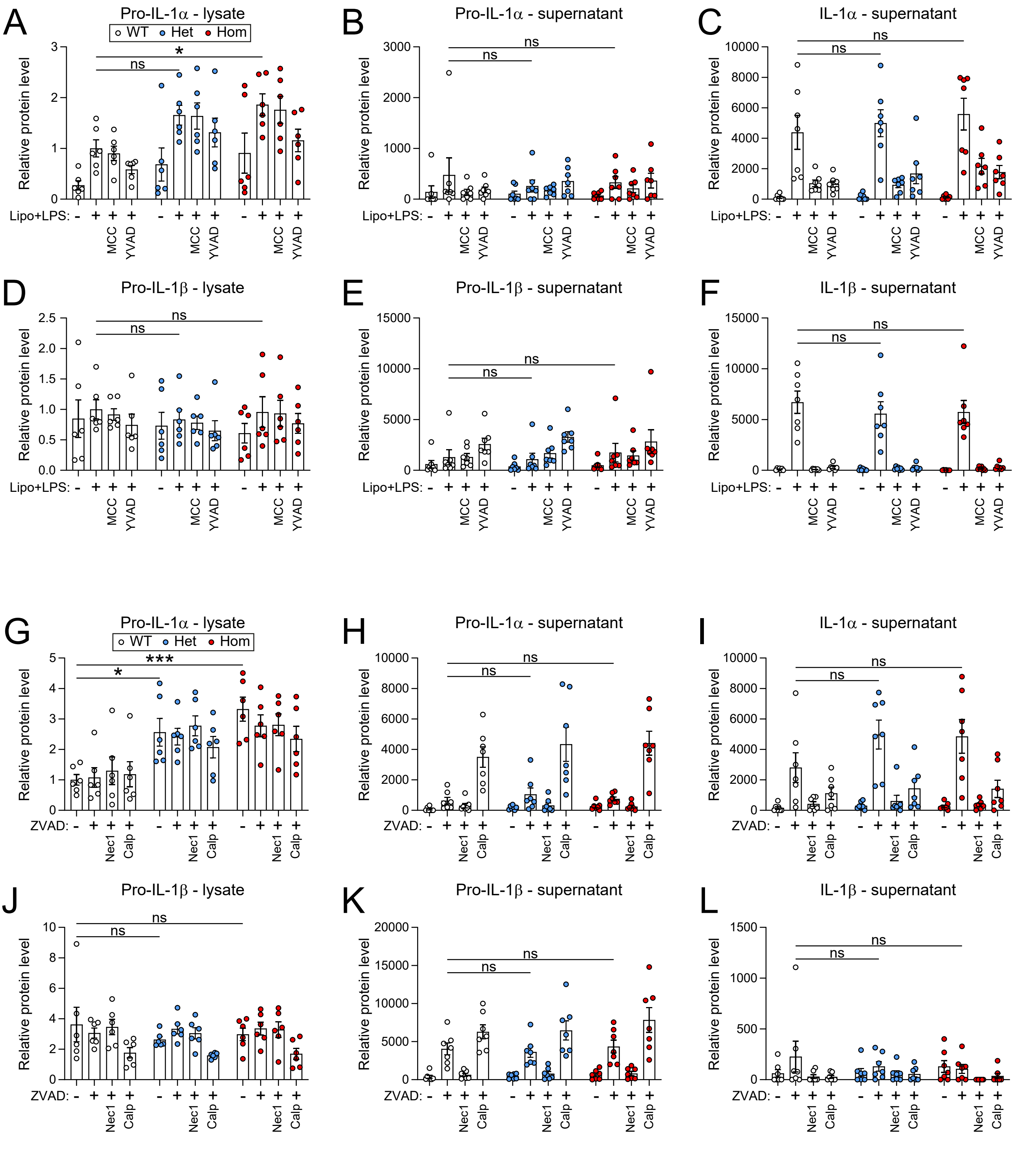


**Supplementary Figure 4. Pro-IL-1α NLS mutation does not negatively affect processing and release of IL-1α in response to non-canonical inflammasome activation or ZVAD-induced necroptosis. (A-F)** Densitometry of western blots from Figure 2H (n=6-7). (A) Relative pro-IL-1α levels were determined in the lysate, and (B) pro-IL-1α and (C) mature IL-1α levels were determined in the supernatant. (D) Relative pro-IL-1β levels were determined in the lysate, and (E) pro-IL-1β and (F) mature IL-1β levels were determined in the supernatant. **(G-L)** Densitometry of western blots from Figure 2L (n=6-7). (G) Relative pro-IL-1α levels were determined in the lysate, and (H) pro-IL-1α and (I) mature IL-1α levels were determined in the supernatant. (J) Relative pro-IL-1β levels were determined in the lysate, and (K) pro-IL-1β and (L) mature IL-1β levels were determined in the supernatant. Data are presented as mean ± SEM. Data were analysed using one-way ANOVA followed by Dunnett's post-hoc test (A,C,F,G,I,K) or Kruskal-Wallis test followed by Dunn's post-hoc test (B,D,E,H,J,L). ***P<0.001; *P<0.05; ns, not significant.

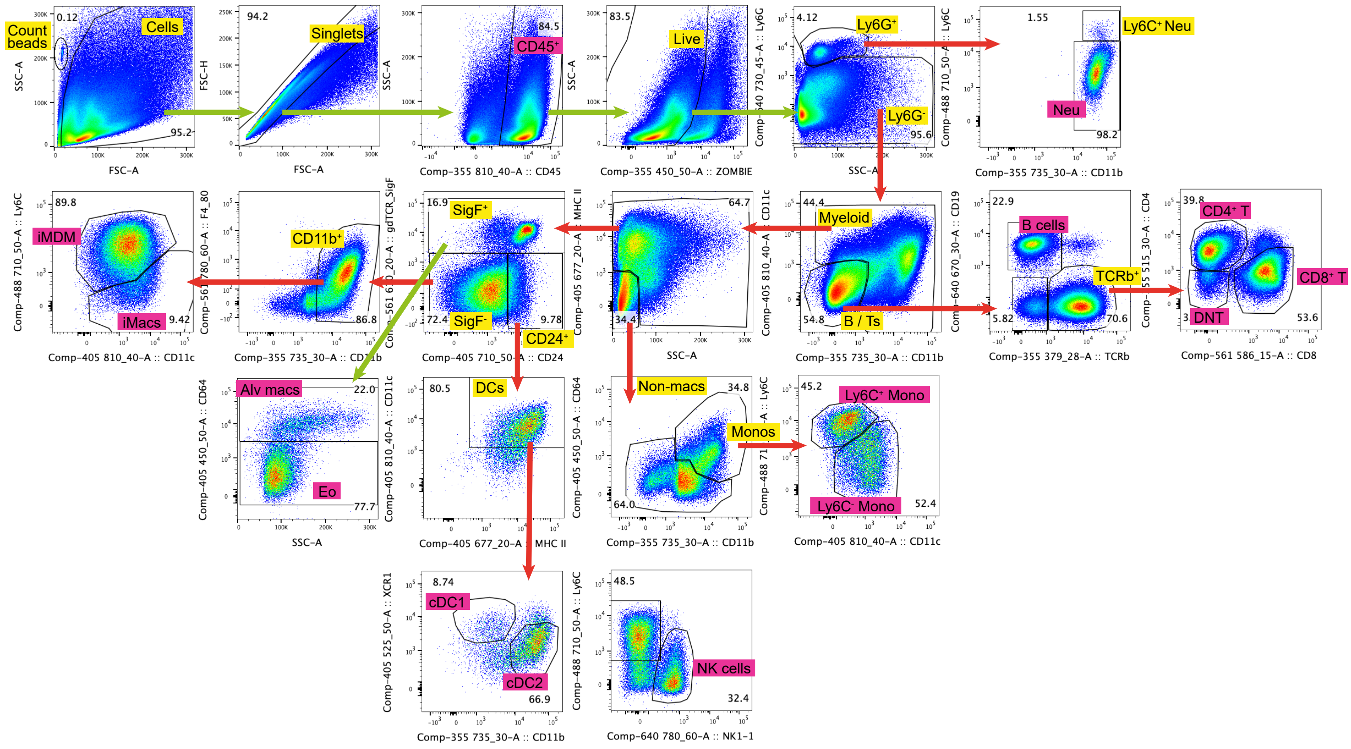


**Supplementary Figure 5. Flow cytometry gating strategy following in vivo IAV infection.** Neu (neutrophils); DNT (CD4$^-$ CD8$^-$ non-classical T cells); Ly6C$^+$ Mono (Ly6C$^+$ classical monocytes); Ly6C$^-$ Mono (Ly6C$^-$ monocytes); NK cells (natural killer cells); cDC1/cDC2 (conventional dendritic cell subtypes 1/2); Eo (eosinophils); Alv macs (alveolar macrophages); iMDM (inflammatory monocyte-derived macrophages); iMacs (interstitial macrophages).

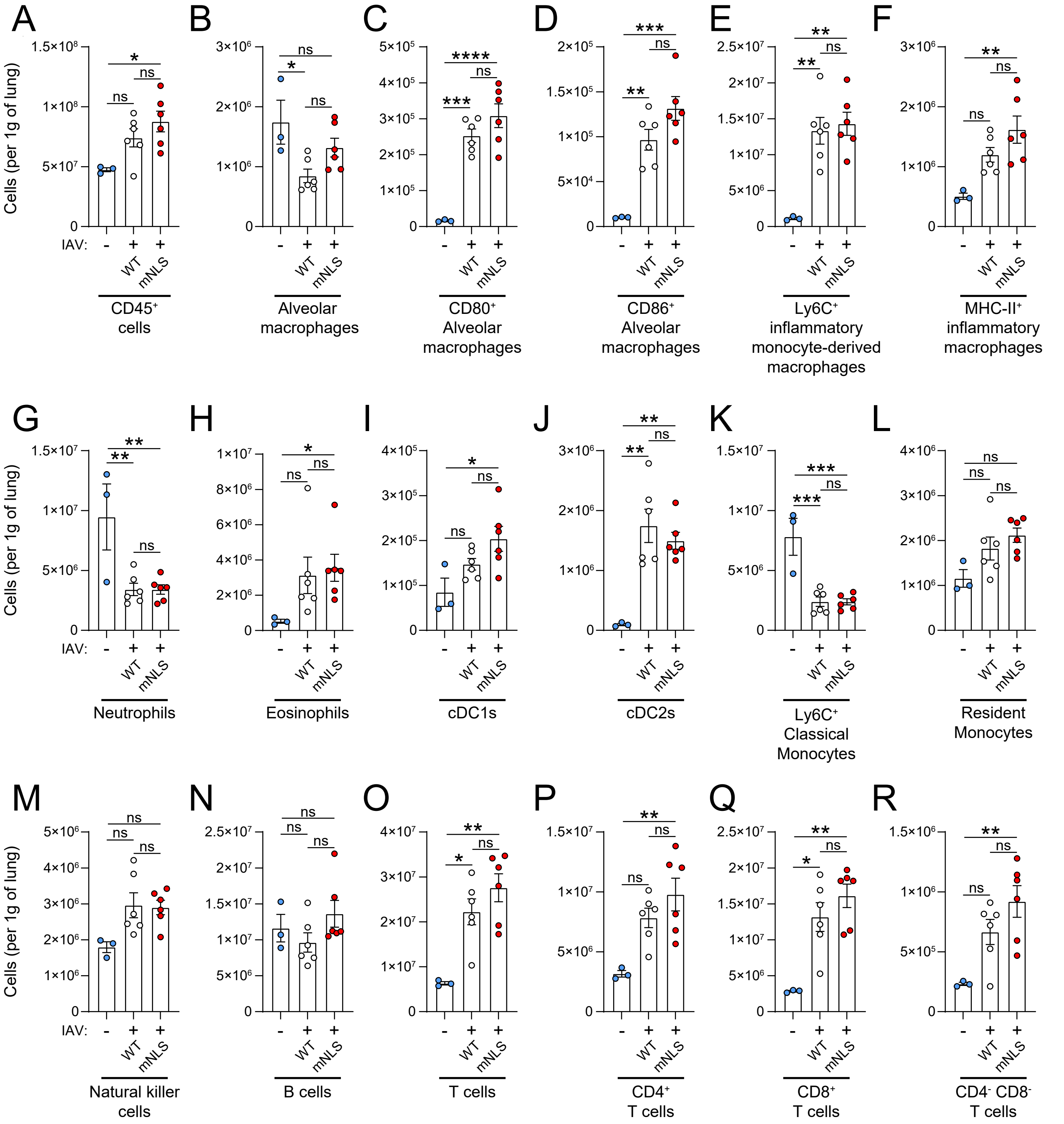


**Supplementary Figure 6. Immune subset populations following IAV infection in WT and mNLS mice.** WT (n=6) or mNLS (n=6) mice were infected intranasally with live influenza strain X31 ($10^3$ PFU in 30 μl of PBS), while heterozygous mice (n=3) were left uninfected (naïve) as a control. **(A-R)** Cell populations were determined using flow cytometry. See Figure 5. Data are presented as mean ± SEM. Data were analysed using one-way ANOVA followed by Sidak's post-hoc test (A-G. I-M, O-R) or Kruskal-Wallis test followed by Dunn's post-hoc test (H,N). ***P<0.0001; ***P<0.001; **P<0.01; *P<0.05; ns, not significant.
